# The Diabetes Gene *Tcf7l2* Organizes Gene Expression in the Liver and Regulates Amino Acid Metabolism

**DOI:** 10.1101/2025.04.03.647067

**Authors:** Joanna Krawczyk, William O’Connor, Pedro Vendramini, Mareike Schell, Kiran J. Biddinger, George Pengo, Tiffany Fougeray, Krishna G. Aragam, Marcia Haigis, Wouter H. Lamers, Linus T. Tsai, Sudha B. Biddinger

**Affiliations:** Division of Endocrinology, Boston Children’s Hospital, Harvard Medical School, Boston MA, USA; Cardiovascular Disease Initiative, Broad Institute of MIT and Harvard, Cambridge MA, USA; Cardiovascular Research Center, Massachusetts General Hospital, Harvard Medical School, Boston MA, USA; Center for Genomic Medicine, Department of Medicine, Massachusetts General Hospital, Harvard Medical School, Boston MA, USA; Department of Cell Biology, Blavatnik Institute, Harvard Medical School, Boston MA, USA; Tytgat Institute for Liver and Intestinal Research, Academic Medical Center, University of Amsterdam, Meibergdreef 69-71, 1105 BK,Amsterdam, The Netherlands; Broad Institute of Harvard and MIT, Cambridge MA, USA; Division of Endocrinology, Diabetes and Metabolism, Beth Israel Deaconess Medical Center, Boston MA, USA; Department of Molecular Metabolism, Harvard T.H. Chan School of Public Health, Boston MA, USA

**Keywords:** Zonation, transcription, diabetes, metabolism

## Abstract

TCF7L2 harbors the strongest genetic association with diabetes identified thus far. However, its function in liver has remained unclear. Here, we find using mice with liver-specific deletion, that *Tcf7l2* plays a central role in maintaining hepatic zonation. That is, in the normal liver, many genes show gradients of expression across the liver lobule; in the absence of *Tcf7l2*, these gradients collapse. One major consequence is the disorganization of glutamine metabolism, with a loss of the glutamine production program, ectopic expression of the glutamine consumption program, and a decrease in glutamine levels. In parallel, metabolomic profiling shows glutamine to be the most significantly decreased metabolite in individuals harboring the rs7903146 variant in *TCF7L2*. Taken together, these data indicate that hepatic TCF7L2 has a secondary role in glycemic control, but a primary role in maintaining transcriptional architecture and glutamine homeostasis.

## Introduction

Genome-wide association studies are performed with the expectation that they will identify genes that are important in human physiology and disease. For type 2 diabetes, the major outcome of these efforts is the identification of *TCF7L2*^1^. TCF7L2 encodes a member of the TCF/LEF family of transcription factors, which bind to β-catenin – a transcriptional regulator that is responsive to Wnt signals^1,2^. In addition to being the first gene found to be associated with type 2 diabetes, *TCF7L2* remains the strongest and most significant association, replicated in multiple cohorts of different ancestries^3^. It appears to be involved in almost 20% of type 2 diabetes cases^4^. Furthermore, *TCF7L2* has been implicated in numerous other diseases associated with type 2 diabetes, ranging from cardiovascular disease to cancer to liver disease^5^. Yet, more than a decade after the identification of TCF7L2 as a diabetes gene, the functions of TCF7L2 remain unclear.

Whole body knockouts of *Tcf7l2* are hypoglycemic and die shortly after birth^6^. Knockout of *Tcf7l2* in β-cells promotes their dysfunction^7^ whereas knockout in adipocytes leads to adipocyte hypertrophy, obesity and insulin resistance^8^. These studies suggest that TCF7L2 may act in multiple tissues to promote diabetes.

The effects of *Tcf7l2* disruption in the liver have been discordant, with some studies reporting hypoglycemia and others hyperglycemia^9–14^. Similarly, both increased and decreased susceptibility to steatosis have been reported^15–17^. These discrepancies may be due, at least in part, to complexities of TCF7L2 function. TCF7L2 is a member of the TCF/LEF family of transcription factors that are often redundant with one another; moreover, both TCF7L2 and β-catenin have multiple additional binding partners^18,19^. Some of the mutations used to disrupt TCF7L2 function lacked the β-catenin interaction domain, but retained the DNA binding domain^14^; others lacked the DNA binding domain but retained the β-catenin interaction domain^12^. Though both mutations would disrupt TCF7L2, their secondary effects on β-catenin and the other members of the TCF/LEF family could differ, making it difficult to determine the unique functions of TCF7L2.

Here, we use mice with targeted deletion of exon 1, with a complete loss of function^20^, to define the specific effects of TCF7L2 in the liver. We find that hepatic *Tcf7l2* has little effect on glucose and lipid metabolism; however, it is essential for the organization of gene expression in the liver. *Tcf7l2* deletion leads to diverse metabolic effects, particularly a decrease in glutamine, which is also observed in individuals with the *TCF7L2* SNP rs7903146.

## Materials and Methods

### Animals

*Tcf7l2^Flox/Flox^* mice^20^ were purchased from Jackson Laboratories (strain code: 031436) and group-housed at 21 °C on a 12 hour light/dark cycle (7 am/7 pm) with free access to food, unless otherwise indicated. At the end of the experiment, mice were killed in the *ad libitum* state at 2 pm EST to collect plasma and tissues. All animal experiments were performed under approval of the Institutional Animal Care and Research Advisory Committee (IACUC) at Boston Children’s Hospital.

*Tcf7l2^Flox/Flox^* littermates were injected retro-orbitally at five to seven weeks of age with 1 x 10^11^ genome copies of AAV8-TBG-GFP (CON, Penn Vector Core, 105535-AAV8) or AAV8-TBG-CRE (TCF7L2 L-KO, Penn Vector Core 107787-AAV8). They were then placed on a Western Diet for twelve weeks (Envigo, TD.88137, irradiated, 42% kcal from fat, 0.2% cholesterol, and 34% sucrose by weight), or continued on a chow diet.

#### Glucose and Lipid Phenotyping

For glucose tolerance testing (GTT), animals were fasted for 14-16 hours overnight, and 2 g D-glucose/ kg body weight was administered intraperitoneally (*i.p.*). For pyruvate tolerance testing, mice were fasted four hours prior to *i.p.* administration of 2 g/kg of sodium pyruvate. For insulin tolerance testing (ITT), mice were fasted four hours prior to administration of a 1 unit/ kg *i.p.* injection of human insulin. For GTT, PTT and ITT, blood glucose levels were monitored via tail nick at 0, 15, 30, 60, 90, and 120 minutes after injection using a glucose meter (Bayer Contour). Values above the limit were assigned the upper limit value of 600 mg/dl. One (control group) or two (L-KO group) mice in each group became hypoglycemic during the ITT and were excluded from the analysis. Additionally, mice were food-deprived for four hours to collect plasma samples in the fasted state using EDTA-coated Eppendorf tubes.

### Plasma analysis

Plasma insulin was measured with an ELISA from CrystalChem (CrystalChem Ultrasensitive Mouse Insulin ELISA). Plasma triglycerides and cholesterol were measured using a colorimetric assay (Infinity Triglycerides, Infinity Cholesterol, Thermo Scientific). Plasma glutamine was measured using a colorimetric assay (Glutamine Assay Kit, Sigma).

### Hepatic Lipids

Hepatic lipids were isolated as previously described^21^. In brief, approx. 80 mg of liver were mechanically homogenized in 50 mM NaCl and extracted with 2:1 chloroform:methanol. The interphase was washed with 50 mM NaCl and 360 mM CaCl_2_ in 50% methanol. The organic extract or positive control lipids (Glycerol Standard, Cholesterol E Standard) were aliquoted and mixed with 10% Triton X-100 (Sigma) in acetone and dried at room temperature overnight. Colorimetric reagents were used as per manufacturers’ recommendations for readout of hepatic lipid levels (Infinity Triglycerides, Infinity Cholesterol, Thermo Scientific; Free Cholesterol E, Wako).

### Bulk gene expression analysis

Total RNA was isolated from frozen liver tissue using TRIZol reagent (Life Technologies). Following isolation, RNA concentration was quantified by a NanoDrop spectrophotometer (Thermo Fisher Scientific) and reverse transcribed to complementary DNA using the High Capacity Reverse cDNA Transcription Kit (Applied Biosystems). Reverse transcriptase quantitative polymerase chain reaction was performed on 25 ng cDNA per well using SYBR green master mix (Applied Biosystems) and 300 nM of each forward and reverse primer (obtained from MilliporeSigma) (see Table 1). Fluorescence was monitored using the Applied Biosystems QuantStudio 6 flex (Thermo Fisher Scientific). Each run was followed by a melt curve (90 °C to 60 °C) for quality control. Samples were analyzed in duplicate, and relative quantification of gene expression levels was performed according to the ΔΔCT method using TATA-box binding protein (*Tbp*) as reference gene. Data were expressed as 2^-ΔΔCT^ and relative to the respective control group, if not stated otherwise.

**Table 1:**
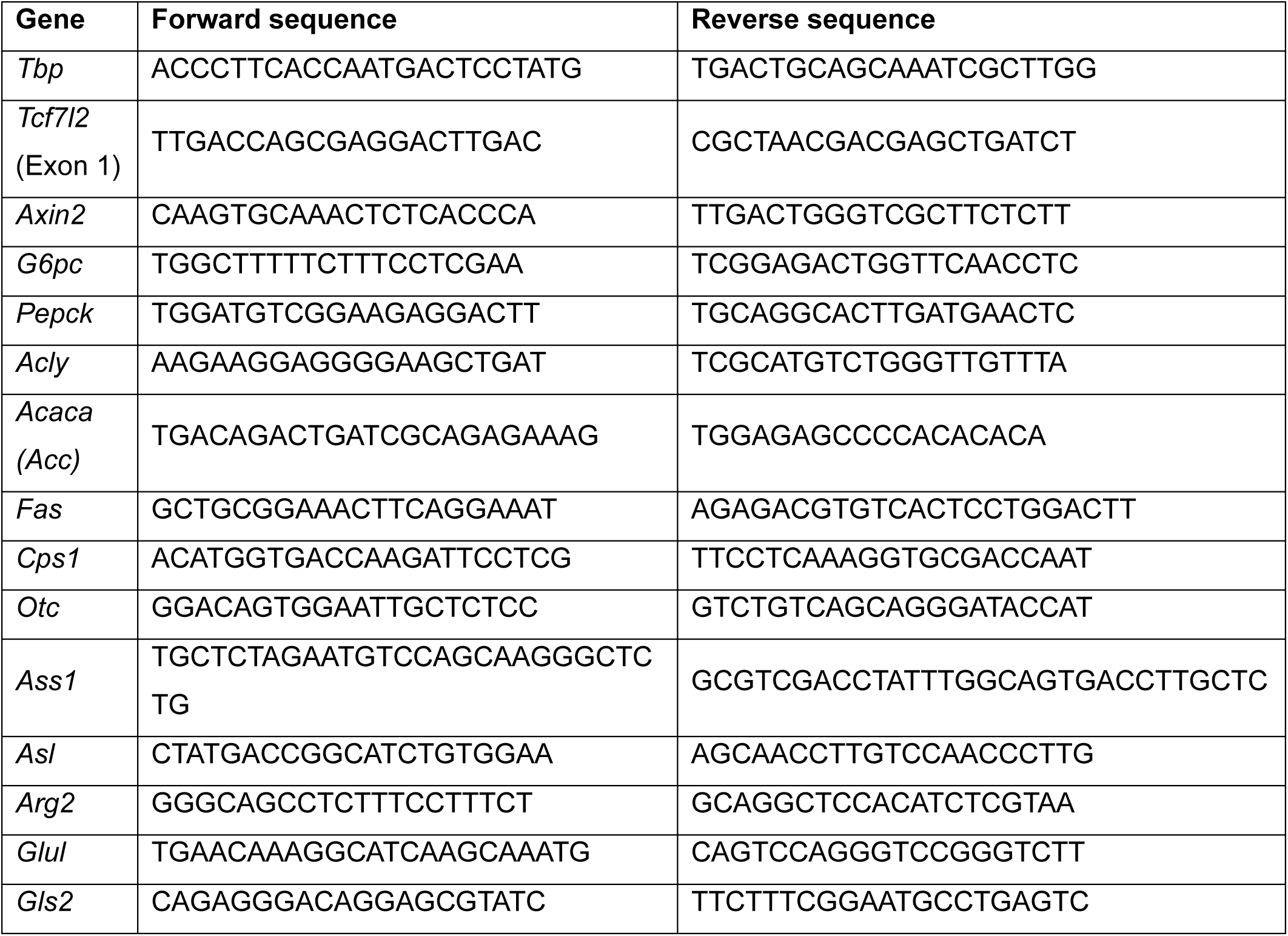
RT-qPCR Primers

### Metabolite profiling and mass spectrometry

#### Extraction of polar metabolites from liver

Metabolites from liver tissue were extracted in 600 µl of -20 °C 60% LC-MS methanol and 400 µl LC-MS chloroform. Samples were vortexed for 15 minutes at 4 °C and centrifuged at 13,300 rpm for 15 min at 4 °C. The supernatant containing polar metabolites was concentrated and dried at 4 °C using a CentriVap SpeedVac (Labconco). The dried extracts were then reconstituted according to tissue weight in 1:1 acetonitrile: water for analysis on the MS (50 µl per mg of tissue to normalize the extraction weight).

#### LC-MS and data analysis

Samples were run on a Vanquish UHPLC system (Thermo Fisher) coupled to a Q-Exactive HF-X mass spectrometer utilizing a HESI probe (Thermo Fisher) in negative ion mode. For the LC, hydrophilic interaction liquid chromatography (HILIC) column was used, specifically a 150 X 2.1 mm iHILIC1-(P) Classic polymeric column equipped with a 2.1 x 20 mm iHILIC1-(P) Classic Guard column (both 5 μm, 200 Å, HILICON AB). Buffer A (20 mM ammonium carbonate in water, with 0.1% ammonium hydroxide) and buffer B (100% acetonitrile) were used. A linear gradient was performed at a flow rate of 0.15 ml/min as follows: 0-23 min linear gradient from 95% B to 5% B; 23-25 min hold at 5% B, to waste from 25-25.5 min gradient to 95% B at 0.20 ml/min, 25.5-32.5 min hold at 95% B, and finally 32.5-33 min 95% B at 0.15 ml/min. The column was held at 25 °C, the column preheater at 30 °C and the autosampler at 4 °C. MS acquisition was performed on full scan mode over a range of 70–1000 m/z and a resolution of 60,000, an AGC target of 1e5 and a maximum injection time of 20 ms with a 5 eV in-source CID. The spray voltage was set to 3 kV, the heated capillary was set at 275 °C and the probe was set at 350 °C. The sheath gas flow rate was 40, the auxiliary gas was set at 15, and the sweep gas flow rate was set to 1. Feature extraction and peak integration were performed using Tracefinder version 4.1 (Thermo Fisher). Metabolites were identified using exact mass with a 5 ppm tolerance and retention time with reference to an in-house library of chemical standards. Peaks were manually curated to exclude low-quality peaks (e.g., noise, multiple peaks) and metabolites with no identified peaks. For statistical and pathway analysis, MetaboAnalyst 6.0 and GraphPad Prism were used with a threshold of -log10(pval)>1.3 (= pval<0.05) and log2(fold change)>|1.0|.

### Immunoblotting

150 mg liver tissue was dounced 24x in a glass tissue grinder set (Corning) with pestle A and B using nuclei EZ prep buffer (Sigma Aldrich, Nuc-101) with added phosphatase inhibitor (Roche). After a five minute incubation on ice, the sample was run through a 100 µm and then a 40 µm filter prior to centrifugation (500 x G for 5 min at 4 °C) and the supernatant was removed (cytosolic fraction). The pellet was washed once in EZ prep buffer and twice with buffer ST (5 mM Tris, 73 mM NaCl, 0.5 mM CaCl_2_, and 10.5 mM MgCl_2_). Nuclei were then spun down, supernatant removed, and nuclear lysis buffer (10 mM HEPES pH 7.9, 0.4 M NaCl, 1 mM sodium EDTA, 1 mM sodium EGTA, 0.5 mM DTT) was added to each sample. Samples were agitated gently at 4°C for one hour, and then centrifuged (13,000 rpm at 4 °C for 10 min). Supernatant was then removed and used for immunoblotting. Protein quantification measurements were performed using the Pierce BCA Protein Assay Kit (Thermo Fisher). Lysates were supplemented with 5X Laemmli sample buffer (10% SDS, 312.5 mM Tris pH 6.8, 0.01% Bromophenol Blue, 50% Glycerol) with freshly added β-mercaptoethanol to 5% then boiled. 10-40 µg of protein was subjected to SDS-PAGE and transferred onto a PVDF membrane at 4 °C (Thermo Scientific). Membranes were incubated for one hour at room temperature 5% milk in 0.1% TBS-T and then overnight at 4 °C with the primary antibody at the concentrations indicated below. The following primary antibodies were used: TCF7L2 (CST2569, 1:1000), and NUP98 (CST2598, 1:1000) from Cell Signaling. Membranes were then incubated with HRP-conjugated secondary antibody (1:15,000 Anti-Rabbit HRP, Cell Signaling). Finally, proteins were detected using SuperSignal West Pico PLUS or DURA Chemiluminescent Substrate (Thermo Scientific).

### Single Nucleus Sequencing

#### Single Nucleus Sequencing Analysis

Nuclei were isolated as previously described^22^. 10X genomics CellRanger Count (v6.1.2)^23^ was used to align FASTQs against the mouse genome (mm39)^24^. Ambient RNA was removed with CellBender (v0.2.0)^25^, and gene-by-cell count matrices were generated using Seurat (v4.3.0.1)^26–30^. scDblFinder (v1.14.0)^31^ was used to identify and remove doublets, and cells with less than 500 detected genes (nFeatures) were removed. Additionally, cells with at least 2% of reads mapping to mitochondrial genes (mt.percent) were discarded. The resulting cells were then normalized using SCTransform^32,33^, and the 3000 most variable features were used for discerning the integration anchors, and the following integration. The first twelve components of principal component analysis were used for finding nearest neighbors, and the Louvain method was used for graph-based clustering (resolution of 0.12). These clusters were identified by known marker genes^34^. T-tests were performed on the expression of each gene in HEP2 versus HEP1 for each genotype. *P* values were corrected using the Benjamini-Hochberg procedure^35^, and log_2_ fold change was plotted for genes with FDR<0.05 and expression in >10% of cells in each genotype. Differential expression was performed for genes between genotypes for each of the two clusters identified as hepatocytes (HEP1, HEP2)^36,37^, and the top 50 genes were subjected to Enrichr analysis^38–41^. Nuclei in clusters identified as hepatocytes were then re-integrated and re-clustered using the top 30 PCs and a resolution of 0.9, resulting in 19 subclusters, four of which were excluded from further analysis due to being distinct from the main body of hepatocytes in the UMAP analysis. The remaining 15 clusters were subjected to diffusion map dimensionality reduction with the first 30 principal components using Destiny (v3.14.0)^42^. The diffusion pseudotime (DPT) was scaled from 1 to 9 and renamed as diffusion pseudodistance (DPD), and lobular location was determined based on previously published pericentral and periportal hepatocyte signatures^43^. First, diffusion pseudodistance values were grouped into 80 bins and a generalized additive model was created for each gene and genotype using gam (v 1.22-2)^44^. The genes were then classified as significantly zonated if *P*<0.05 based on a parametric ANOVA of the model in either or both genotypes. Diffusion pseudodistance values were then grouped into 8 bins for each genotype. Hierarchical clustering was performed on all bins from both genotypes for each significantly zonated gene, and four groups were created (hclust, cutree, stats v4.3.1). Pathway and transcription factor analysis was performed on each cluster using Enrichr^38–41^. Overrepresentation analysis was also performed using a hypergeometric test on each cluster and genes known to be regulated by β-catenin^45,46^. *P* values were adjusted using the Benjamini Hochberg procedure^35^. The DPT 8 bins for each genotype were also used to show the expression of glutamine and glutamate genes, and hierarchical clustering was once again performed^41,47^.

### Immunofluorescence and smFISH

#### Sectioning

The right posterior lobe of the liver was fixed in 10% neutral buffered formalin at 4 °C overnight with shaking. Samples were washed four times with shaking in phosphate-buffered saline (PBS), then stored in 70% ethanol until further processing. Paraffin embedding and cutting (5 µm sections), as well as hematoxylin and eosin staining, were performed by the Beth Israel Deaconess Histology Core.

#### Single Molecule In Situ Hybridization

*In situ* hybridization was performed on 5 μm formalin fixed, paraffin embedded (FFPE) sections using RNAscope multiplex fluorescent V2 assay (323110, Advanced Cell Diagnostics, Inc). The following probes were used: Mm-Cps1-C2 (437201-C2); Mm-Gls2-C3 (449281-C3), and Mm-Glul-C1 (426231). Tissue pretreatment was performed according to the manufacturer’s instructions for FFPE murine liver tissue. Hybridization, downstream amplification steps, and DAPI staining/mounting were performed according to the manufacturer’s protocol (UM 323100). Transcripts were labeled with PerkinElmer TSA Plus fluorophores. Glul-C1 probe was detected using TSA Plus fluorescein (NEL741E001KT); Cps1-C2 probe was detected with TSA Plus Cyanine 3 (NEL744E001KT), and Gls2-C3 probe was detected with TSA Plus Cyanine 5 (NEL745E001KT).

#### Image Collection and Downstream analyses

Images were collected at 20X with a Nikon upright Eclipse90i microscope; five Z-stacks with steps of 1 μm were combined to form the focused image, and downstream adjustments were performed in ImageJ. In all cases, images were collected using the same exposure time across samples, and downstream adjustments were performed equally across all experimental groups. One to five images were collected per mouse. Quantification was done using custom Python scripts^48^ (please see: *Data availability*). Transcripts were detected by the Big-FISH Python package^49^. Central veins were identified by the marker gene *Glul* and/or absence of a bile duct. Central and portal veins were outlined in CellProfiler^50^ Transcripts were assigned to 10 bins based on distance from the central vein relative to the sum of distances to the central and portal veins. The counts in each bin were normalized by the area of the bin in pixels, and the percentage of total counts in each bin was calculated. Values were then averaged by mouse.

### Human genetics studies

The UK Biobank is a large population-based biobank consisting of extensive genetic and phenotypic data for 502,629 participants^51^. Detailed baseline plasma metabolic biomarker quantification was performed in 118,461 of these patients^52,53^. 10,442 individuals were excluded based upon sample-level reliability metrics and availability of phenotypic data. Metabolite or disease associations with the rs7903146 variant in the TCF7L2 gene were tested in the 108,019 remaining individuals using linear or logistic regression adjusted for participant age at baseline assessment, sex, and the top ten principal components of genetic ancestry. Supplemental Table 4 lists the phenotypes assayed. Informed consent was obtained for all UK Biobank study participants, and analysis was approved by the Mass General Brigham Health Care Institutional Review Board (under protocol 2013P001840; UK Biobank application 7089).

### Statistical Analysis

Data are represented as mean +/-SEM, unless stated otherwise. For comparisons between two groups, a Students t-test was performed with \**P* < 0.05, \*\**P* < 0.01, \*\*\**P* < 0.001, \*\*\*\**P* < 0.0001. All statistical analysis was performed in GraphPad Prism (Version 10) and R (v4.3.1).

### 2.11. Data Availability

All data are available in the main text or the supplementary materials (https://www.dropbox.com/scl/fo/da7g0krtpawm4tc7x7bt1/AIjZa9Ml8Tb-5lDGp7XvwH8?rlkey=x4xbeqlx20n1ci4q8bw4qt1ns&st=p0uv3xao&dl=0), with the exception of the Single Nucleus Sequencing data, which will be submitted to GEO upon acceptance of manuscript. Custom R and Python scripts can be found at https://github.com/WO-Connor/TCF_Knockout/.

## Results

Mice harboring floxed alleles of exon 1 of the *Tcf7l2* locus were injected with adeno-associated virus encoding Cre recombinase under the *Tbg* promoter to generate mice with Liver specific Knockout (L-KO) of *Tcf7l2*. For controls (CON), we used littermates injected with adeno-associated virus in which green fluorescent protein was substituted for the *Cre* recombinase.

In the absence of any overt phenotype on the chow diet (Supplemental Figure 1), we subjected mice to the stress of a Western diet (34% sucrose by weight; 42% fat; 0.2% cholesterol). Figure 1A shows an immunoblot of nuclear liver extracts. Using an antibody against an epitope close to the HMG-box DNA binding domain^54^, we observed two major bands at 58 and 79 kDa in the control livers, as previously reported^15^. These were undetectable in the L-KO mice. We found no overt phenotype on this diet either (Figure 1B-1L). Body weight was unchanged. Plasma glucose was measured in the *ad libitum* fed state (data not shown); in the four hour fasted state; during a glucose tolerance test; during a pyruvate tolerance test; and during an insulin tolerance test. All were normal. Plasma and hepatic lipid levels also remained similar to controls, though there was a trend towards reduced hepatic triglycerides. In parallel, expression of the gluconeogenic and lipogenic genes, assayed on whole liver homogenates using QPCR, was similar between genotypes.

**Figure 1:**
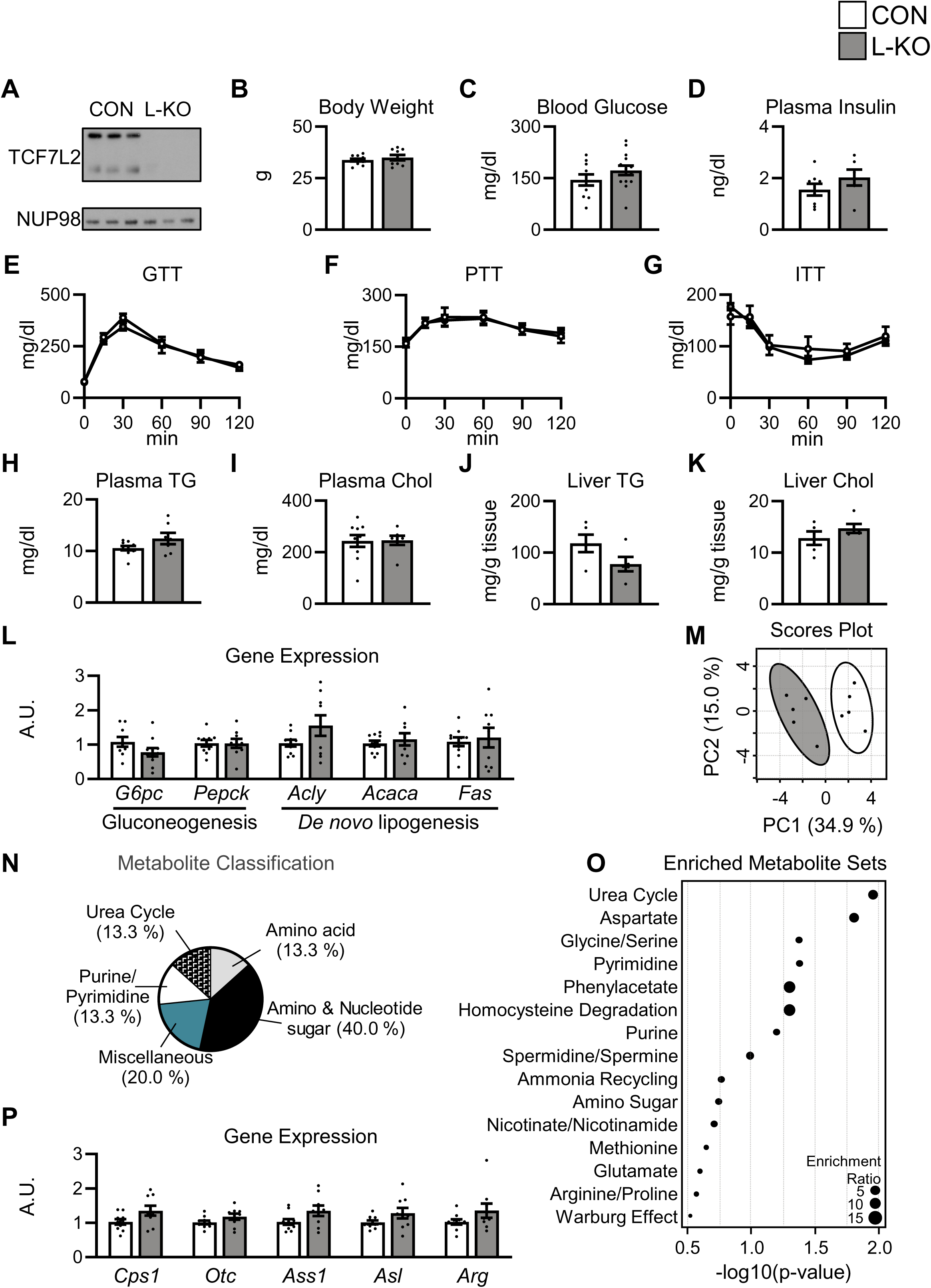
Disruption of hepatic *Tcf7l2* leads to widespread metabolite changes. Six to eight week old male *Tcf7l2^Flox/Flox^* mice were injected with adeno-associated virus encoding either GFP (CON) or Cre (L-KO) and placed on a western diet for twelve weeks. (**A**) Western blot of liver nuclear fractions. (**B**) Final body weight. (**C, D**) Four hours fasting blood glucose (**C**) and (**D**) plasma insulin. (**E**) Blood glucose levels during glucose tolerance test (GTT), (**F**) pyruvate tolerance test (PTT), and (**G**) insulin tolerance test (ITT). Four hours fasting (**H**) plasma triglycerides (TG) and (**I**) cholesterol (Chol). (**J**) Liver triglycerides (TG) and (**K**) cholesterol (Chol). (**L**) Liver QPCR analysis. (**M-O**) Liver metabolite profiling. (**M**) Principal Component Analysis (PCA). (**N**) Significantly altered liver metabolites (-log10(pval)>1.3). (**O**) Metabolite Set Enrichment Analysis (MSEA). (**P**) Liver QPCR analysis. Data are presented as the mean ± SEM; n=5-14/group. A.U., arbitrary units.

We next performed metabolomics analysis, measuring 183 metabolites in the livers of control and L-KO mice. Interestingly, principal component analysis revealed a clear separation by genotype, indicating significant metabolic derangements in the *Tcf7l2* knockout mice on both the chow (Supplemental Figure 1M-1O) and Western diet (Figure 1M-1O). Most striking were the changes in amino acid metabolism, which accounted for more than one third of the significantly altered metabolites. In parallel, pathway analysis showed the urea cycle, which is required for the catabolism of amino acids, to be the most dysregulated pathway. Yet, QPCR on whole liver homogenates did not reveal robust changes in urea cycle gene expression, raising the question of how the transcription factor TCF7L2 alters metabolism (Figure 1P, Supplemental Figure 1P).

Though the liver is often considered a homogenous organ, it has a complex microarchitecture, with millions of lobules. Each lobule is anatomically defined by a central vein, surrounded by concentric layers of hepatocytes, with portal triads located at the periphery (Figure 2A). Those hepatocytes closest to the central vein (i.e., pericentral hepatocytes), and those closer to the portal vein (i.e., periportal hepatocytes) differ in their transcriptional profiles^43,55^. Importantly, a key binding partner of TCF7L2, β-catenin, is activated only in the pericentral hepatocytes, suggesting that TCF7L2 might have distinct effects in those hepatocytes.

**Figure 2:**
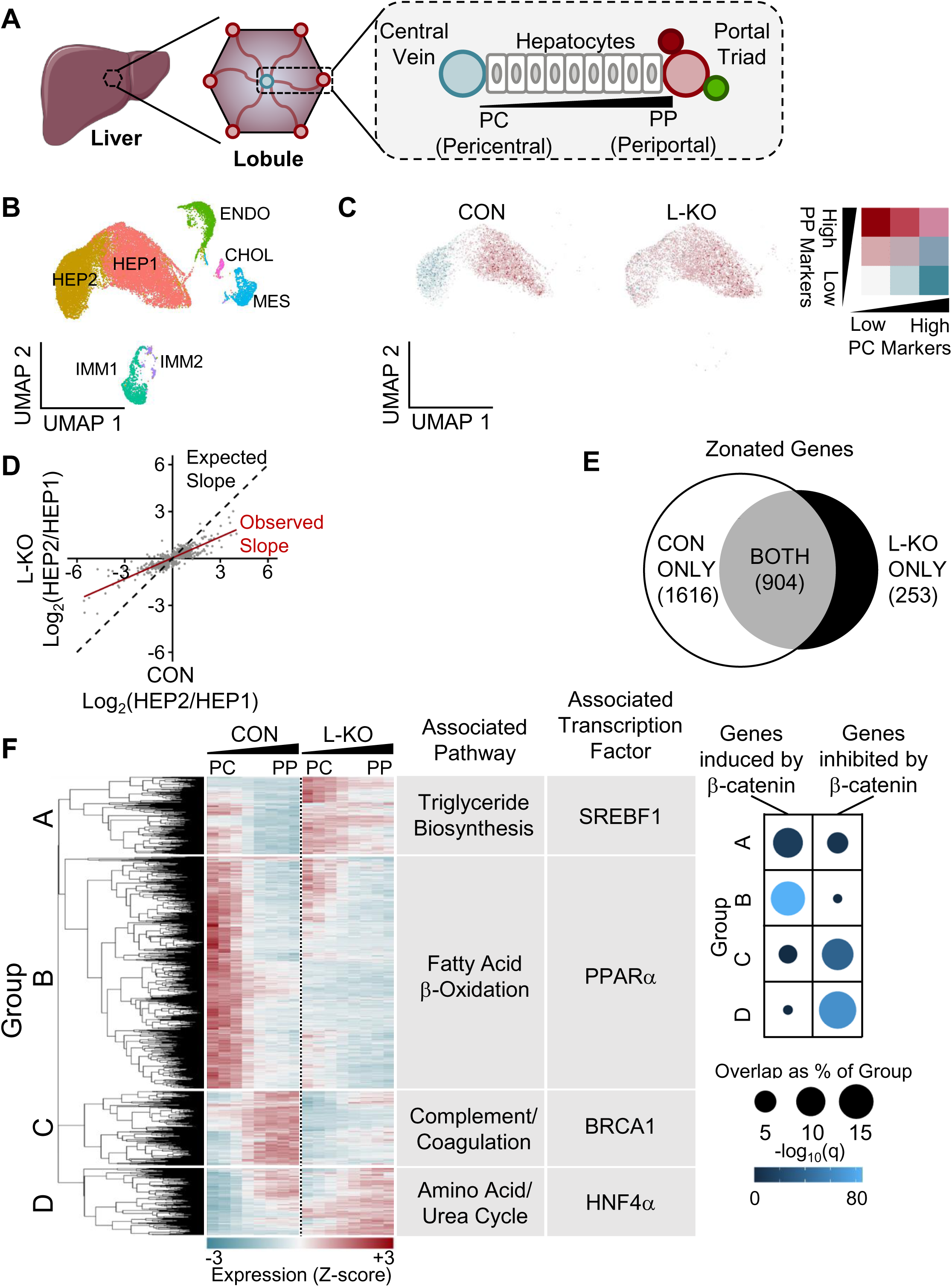
*Tcf7l2* is required for hepatic zonation. (**A**) Anatomy of the liver lobule. (**B-F**) Single nucleus sequencing was performed on the livers of five to six week old *Tcf7l2^Flox/Flox^* mice injected with adeno-associated virus encoding either GFP (CON) or Cre (L-KO) and placed on a western diet for 12 weeks. (**B**) UMAP projection of 24175 nuclei from the livers of CON and L-KO (HEP1, hepatocyte 1; HEP2, hepatocyte 2; ENDO, endothelial cells; CHOL, cholangiocytes; MES, mesenchymal cells; IMM1, immune 1; IMM2, immune 2). (**C**) Hepatocyte clusters (n=18242 nuclei) visualized by periportal (PP) and pericentral (PC) marker expression. (**D**) Distribution of gene expression among the two hepatocyte clusters in CON livers versus L-KO livers. (**E**) Number of significantly zonated genes by genotype. (**F**) Analysis of zonated genes: unsupervised clustering; top associated transcription factor and pathway identified by Enrichr^38–40,65,66^; and overrepresentation^45^ of genes previously shown to be induced or suppressed in response to β-catenin^46^. Full Enrichr results are shown in Supp. Table 3.

We therefore performed single nucleus sequencing on the livers of five control mice and five *Tcf7l2* knockout mice. 29,637 nuclei were identified, of which 24,175 passed quality control. UMAP analysis was performed (Figure 2B, Supplemental Figure 2A-2D). Both control and KO livers contributed similar proportions of cells to each cluster (Supplemental Figure 2E). The clusters were presumptively identified by the presence of marker genes, with HEP1 and HEP2 as hepatocyte clusters^34^. On the one hand, in control mice, HEP1 showed high expression of periportal markers, but low expression of pericentral markers; conversely HEP2 showed high expression of pericentral markers, but low expression of periportal markers^43^ (Figure 2C). These patterns identify HEP1 and HEP2 as the periportal and pericentral nuclei, respectively, and highlight the robust transcriptional differences between them. In L-KO mice, on the other hand, pericentral markers were low in both HEP1 and HEP2, and periportal markers were high in both HEP1 and HEP2. These data show a disruption in the normal zonation profiles and suggest that the transcriptional distinctions between the two hepatocyte clusters were attenuated in the knockout mice.

To determine the effects TCF7L2 on global gene distribution, we calculated the ratio of expression in HEP2 versus HEP1 for differentially expressed genes in each genotype, expressed it as a log fold change, and then plotted the value for the knockout livers against that of the control livers (Figure 2D). The resulting slope was less than 1, indicating that the gene expression differences between the two hepatocyte clusters observed in control mice was, on the whole, attenuated in L-KO mice. These data confirm that the differences between the two clusters is diminished in the absence of TCF7L2.

To examine, with greater granularity, the effects of TCF7L2 on zonation, we performed diffusion pseudodistance analysis. Here, the distance of each nucleus from the central vein was inferred from diffusion map dimensionality reduction^43^, enabling us to assess gene expression quantitatively across the lobule. This analysis showed 1616 transcripts were significantly zonated in control mice—that is, they showed a non-uniform distribution across the lobule; only 904 of these transcripts retained zonation in the knockout mice (Figure. 2E). Moreover, 253 genes became significantly zonated in knockout mice.

Unsupervised clustering of the genes that were significantly zonated in either genotype produced four groups. These groups were found to be associated with distinct metabolic processes and transcription factors^38–40^ (Figure 2F). Group A contained genes normally restricted to the pericentral region of control mice but expressed diffusely in L-KO mice. This group was enriched for the genes of triglyceride biosynthesis and the SREBF1 gene signature. Group B contained genes normally enriched in the pericentral region, but reduced upon deletion of *Tcf7l2*. This group was enriched for the genes of fatty acid β-oxidation and the PPARα gene signature. Group C contained genes normally enriched in the periportal region, but reduced upon deletion of *Tcf7l2*. This group was enriched in the genes involved in complement and coagulation and the BRCA1 gene signature. Finally, Group D contained genes normally restricted to the periportal region, that diffusely expressed in L-KO mice. This group was enriched in the genes of amino acid and urea cycle metabolism and the HNF4α gene signature.

To dissect the role of β-catenin, we leveraged previous work that used mice with loss and gain of function of β-catenin in the liver to define the genes induced and inhibited by β-catenin^46^. We found significant overlap between the β-catenin targets and those disrupted by the deletion of *Tcf7l2*. In particular, Group B was highly enriched in genes up regulated by β-catenin, and Group D was highly enriched in genes down regulated by β-catenin^46^.

To better understand the physiological differences between L-KO and control mice we performed differential gene expression analysis within each zone^38–41^, focusing on the pericentral zone (HEP2), which appeared to have the greatest changes (Figure 3A, Supplemental Figure 3A). Among the genes in the pericentral region that were reduced by *Tcf7l2* deletion, the most overrepresented gene set was TCF/LEF signaling, consistent with the loss of Wnt signaling in these mice. The second most overrepresented gene set was glutamate/glutamine metabolism. This was particularly interesting to us given the change in amino acid and urea cycle metabolism observed by metabolomics (Figure 1N, O).

**Figure 3:**
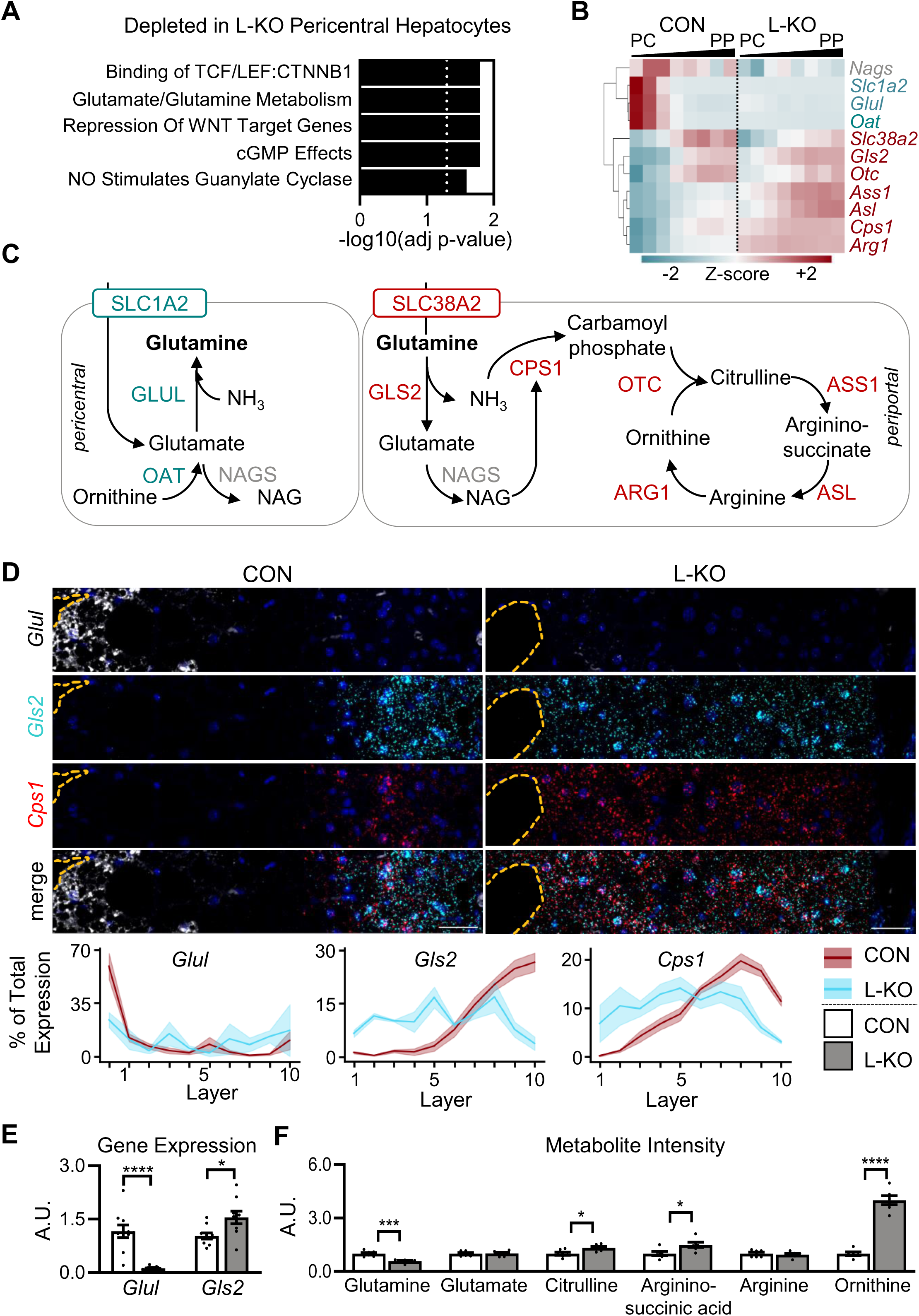
Disruption of hepatic *Tcf7l2* results in suppressed pericentral glutamine production. Five to eight week old male *Tcf7l2^Flox/Flox^* mice were injected with adeno-associated virus encoding either GFP (CON) or Cre (L-KO) and placed on a western diet for 12 weeks. (**A**) Genes that were differentially expressed between pericentral hepatocytes (HEP2) from CON and L-KO livers by snSEQ were subjected to Enrichr analysis as described in methods. Five most significant gene sets are shown. Dotted line marks significance threshold of -log10(adj p-value)>1.3. (**B**) Glutamine metabolism genes were subjected to unsupervised hierarchical clustering. (**C**) Schematic. (**D**) Representative images of smFISH and quantification. (**E**) Liver QPCR analysis. (**F**) Liver metabolites. Data are presented as the mean ± SEM; n=4-8/group. *P* values were determined by Student’s *t*-test; \**P* < 0.05, \*\*\**P* < 0.001, \*\*\*\**P* < 0.0001. Central vein highlighted by dashed orange line; scale bar = 40 µm. A.U., arbitrary units.

Unsupervised clustering of the genes involved in glutamate and glutamine metabolism^41,47^ revealed groups of genes with distinct patterns of gene expression that, interestingly, correlated with not only location, but function (Figure 3B, 3C). One group was enriched in the pericentral nuclei of control mice, and lost in the L-KO mice. It contained genes involved in the production of glutamine (*Slc1a2*, *Glul*, and *Oat*). Another group of genes was enriched in the periportal nuclei of control mice; in the L-KO mice, their expression was increased and more diffuse. It contained genes involved in the consumption of glutamine (*Slc38a2*, *Gls2*, *Otc*, *Ass1*, *Asl*, *Cps1*, *Arg1*).

To confirm these changes, we performed single molecule *in situ* hybridization (smFISH). In control livers, the glutamine synthesis gene, *Glul*, was clearly expressed in the first few layers of hepatocytes surrounding the central vein; in L-KO mice, it was undetectable in any hepatocyte. In contrast, the glutamine consumption gene *Gls2*, as well as the urea cycle enzyme *Cps1*, were restricted to the periportal region in control livers but expanded in the knockout livers, mirroring the snSEQ results. Bulk levels of gene expression, measured by QPCR of liver homogenates, which represent an average gene expression showed L-KO mice to have decreased *Glul* and increased *GIs2* (Figure 3E), despite normal levels of *Cps1* (Figure 1P).

In parallel with these gene expression changes, glutamine levels were significantly reduced in the livers of L-KO mice, whereas the urea cycle intermediates, citrulline, arginino-succinate and ornithine were significantly increased (Figure 3F). Glutamine levels were also reduced 25% in the plasma of L-KO mice (Supplemental Figure 3C).

These studies in mice showed that hepatic *Tcf7l2* had profound effects on amino acid metabolism. To correlate these changes in mice with those in humans, we turned to the UK Biobank^51^. This resource contains both genetic data and baseline plasma measurements of 168 biomarkers, including amino acids, fatty acids, glycolysis metabolites, ketones, inflammatory markers, lipids and lipoprotein measurements (Supplemental Table 4)^52^. Among the patients phenotyped for these metabolites, we identified 54,649 individuals with the CC genotype, 44,178 with the CT genotype, and 9,192 with the TT genotype for the rs7903146 variant in *TCF7L2*, the strongest GWAS signal for type 2 diabetes^1^. These groups were not significantly different by age, but showed slight male predominance. As expected, the SNP was positively associated with type 2 diabetes (odds ratio per T allele, 1.352 [95% CI, 1.298-1.408], p=1.89e-168). The metabolite showing the strongest association was glucose, which was positively associated with the T allele (effect estimate per T allele, 0.055 mmol/l [0.045-0.066], p=1.44e-47). The metabolite showing the next strongest association was glutamine (effect estimate per T allele, -2.137 µmol/l [-2.874--1.399], p=1.37e-08), which was negatively associated with the T allele. The association with glutamine remained significant even after adjusting for diabetes status, as well as age and sex (p=2.79e-06).

## Discussion

Mice with liver-specific knockout of *Tcf7l2* show zone-specific disruptions in gene expression. In particular, the genes for glutamine synthesis, which are normally located pericentrally, are lost, and the genes for glutamine degradation/urea cycle, which are normally periportal, are increased. In parallel, glutamine is decreased in the livers and plasma of knockout mice and the plasma of individuals harboring the rs7903146 variant in *TCF7L2*. Thus, these studies reveal a critical role for TCF7L2 in organizing gene expression in the liver and maintaining metabolic homeostasis.

Previous studies *in vitro* and using bulk liver homogenates have shown a variety of direct and indirect interactions of TCF7L2 with other transcription factors^10,17,18,46^. Here, we find that many of these interactions are zone dependent. The most important zone-specific regulator of TCF7L2 is likely β-catenin. β-catenin can only enter the nucleus, interact with TCF7L2, and regulate gene expression in the pericentral hepatocytes^56^. This is due to at least two factors. First, the WNT ligands that activate β-catenin--particularly WNT2 and WNT9b, which are required for zonation^56^ --are expressed in the central vein endothelial cells and have a very limited signaling radius, potentially due to their lipidation and limited diffusion capability^12,57^. Second, WNT signaling is suppressed in the periportal region due to expression of Wnt antagonists^58,59^. In addition, *Tcf7l1*, a potential repressor of TCF7L2, has been reported recently to be periportally enriched^60,61^. Consequently, TCF7L2 can only activate β-catenin targets in the pericentral hepatocytes.

Chromatin immunoprecipitation studies in the livers of mice with mutations in β-catenin have elegantly shown that TCF7L2 binds to a distinct set of promoters in the presence versus absence of β-catenin^46^. Our data are consistent with this, and suggest a model in which TCF7L2 acts, on the one hand, in the pericentral region as an effector of β-catenin to activate (Group B, Figure 2F) and repress (Group D, Figure 2F) its targets. On the other hand, it appears to act primarily in the periportal region to activate (Group C, Figure 2F) and repress (Group A, Figure 2F) a distinct set of targets.

At the physiological level, hepatic *Tcf7l2* deletion, surprisingly, does not affect plasma glucose levels, which were normal under every condition tested; in parallel, both the bulk levels and the distribution of the gluconeogenic genes are similar to controls (Figure 1L, Supplemental Figure 1L, Supplemental Figure 3B). Plasma and hepatic lipids are also unaffected by *Tcf7l2* deletion (Figure 1H-1K, Supplemental Figure 1H-1K). Nonetheless, in the pericentral hepatocytes of L-KO mice, *de novo* lipogenesis genes are increased, and fatty acid oxidation genes are decreased (Figure 2F).

The metabolism of amino acids, however, is markedly dysregulated both at the transcript and metabolite levels in mice with hepatic deletion of *Tcf7l2*. On the one hand, expression of glutamine degrading enzyme *Gls2* is no longer confined to the periportal hepatocytes and bulk transcript levels are increased. Similarly, the urea cycle genes, which are required for the disposal of the ammonia generated by glutamine catabolism, are also increased, as are several metabolites in the urea cycle. On the other hand, the genes for glutamine production, particularly *Glul*, normally expressed in the pericentral hepatocytes, are lost. Consequently, *Tcf7l2* L-KO mice show a 42%- and 25% decrease in glutamine in their livers and plasma (Figure 3F, Supplemental Fig 3C). Importantly, the role of hepatic TCF7L2 is not sex-specific, as female *Tcf7l2* L-KO mice showed marked derangements in the expression and distribution of the enzymes of glutamine metabolism, but were normoglycemic (Supplemental Figure 4).

In humans, the rs7903146 variant, which presumably affects TCF7L2 in multiple tissues, is associated with reduced levels of plasma glutamine; indeed, the association with glutamine was second only to the association with glucose among the metabolites tested (Figure 4B).

**Figure 4:**
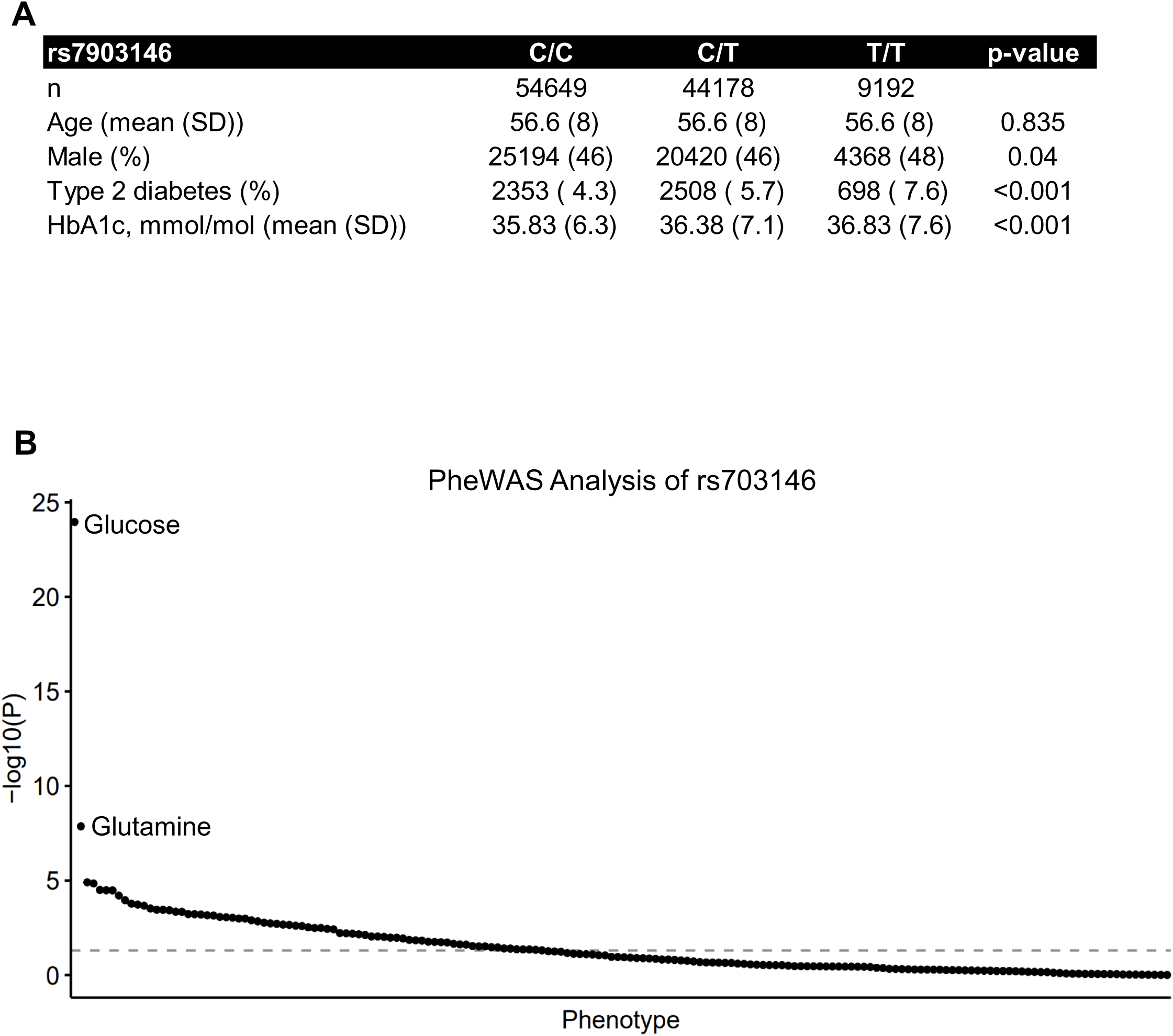
Glutamine associates with TCF7L2 rs7903146 variant. Human phenotypic profiling of TCF7L2 rs7903146 variant within UK Biobank study participants. (**A**) Characteristics of 108,019 UK Biobank participants stratified by rs7903146 carrier status. Values are presented as either number (%) or mean (SD). Nominal p-values calculated as either Chi-squared test for categorical variables or ANOVA for continuous variables. (**B**) Association p-values for rs7903146 carrier status and 168 NMR-derived biomarkers. -log10(p-values) were calculated using linear regression models. Dashed gray line shows Bonferroni-corrected threshold for significance (p = 2.97e-04).

## Conclusions

Taken together, these data suggest that *Tcf7l2* does not act in the liver to directly impair glucose metabolism and that the major effect of *Tcf7l2* must therefore be extrahepatic, potentially impairing β-cell or adipose function. Nonetheless, *Tcf7l2* has a major effect on the organization of transcription in the liver, and widespread metabolic effects. In particular, we find glutamine metabolism is disrupted. Importantly, this defect in glutamine may promote metabolic dysfunction indirectly, as low glutamine levels can impair beta-cell function^62,63^, and glutamine supplementation in humans appears to have beneficial metabolic effects^64^.

## Financial support

National Institutes of Health, T32 training grant DK007260 (JK)

Harvard Digestive Disease Center, P30 DK034854 (Pilot and Feasibility Award to SBB, MH)

American Heart Association, 862032 (KGA)

National Institutes of Health, 5R01DK125898 (SBB) and 1K08HL153937 (KGA)

Deutsche Forschungsgemeinschaft (DFG, German Research Foundation) – Projektnummer 540879521 (MS)

## Acknowledgements

We thank all members of the Biddinger Lab for suggestions and experimental support. We also thank the Harvard Digestive Disease Center Histology Core (NIH P30DK034854), the Boston Children’s Hospital Viral Core (NIH 5P30EY012196), the Boston Nutrition Obesity Research Center Functional Genomics and Bioinformatics Core (NIH P30DK046200).

## Abbrevations

Acaca: Acetyl-CoA carboxylase
HEP: Hepatocytes
Acly: ATP citrate lyase
HNF4α: Hepatocyte nuclear factor 4 α
Arg: Arginase
IMM: Immune cells
Asl: Argininosuccinate lyase
ITT: Insulin tolerance test
Ass1: Argininosuccinate synthase
LC-MS: Liquid chromatography–mass spectrometry
BRCA1: Breast cancer type 1 susceptibility protein
L-KO: Liver knockout
CHOL: Cholangiocytes
MES: Mesenchymal cells
Chol: Cholesterol
Nags: N-Acetylglutamate synthase
CON: Control
Otc: Ornithine transcarbamylase
Cps1: Carbamoyl phosphate synthetase
PBS: Phosphate buffered saline
Cre: Cre recombinase pericentral
DNA: Deoxyribonucleic acid
Pck1/Pepck: Phosphoenolpyruvate carboxykinase
DPD: Diffusion pseudodistance
PP: periportal
DPT: Diffusion pseudotime
PPARα: Peroxisome proliferator-activated receptor α
ELISA: Enzyme-linked immunosorbent assay
PTT: Pyruvate tolerance test
ENDO: Endothelial cells
QPCR: Quantitative Real-time polymerase chain reaction
Fas: Fatty acid synthase
RNA: Ribonucleic acid
G6pc: Glucose-6-phosphatase
smFISH: Single molecule Fluorescence *in situ* hybridization
GFP: Green fluorescent protein
snSEQ: Single cell sequencing
Gls2: Glutaminase 2
SREBF1: Sterol regulatory element-binding transcription factor 1
Glul/Gs: Glutamine synthetase
TG: Triglyceride
GTT: Glucose tolerance test

---

**Suppl. Figure 1:**
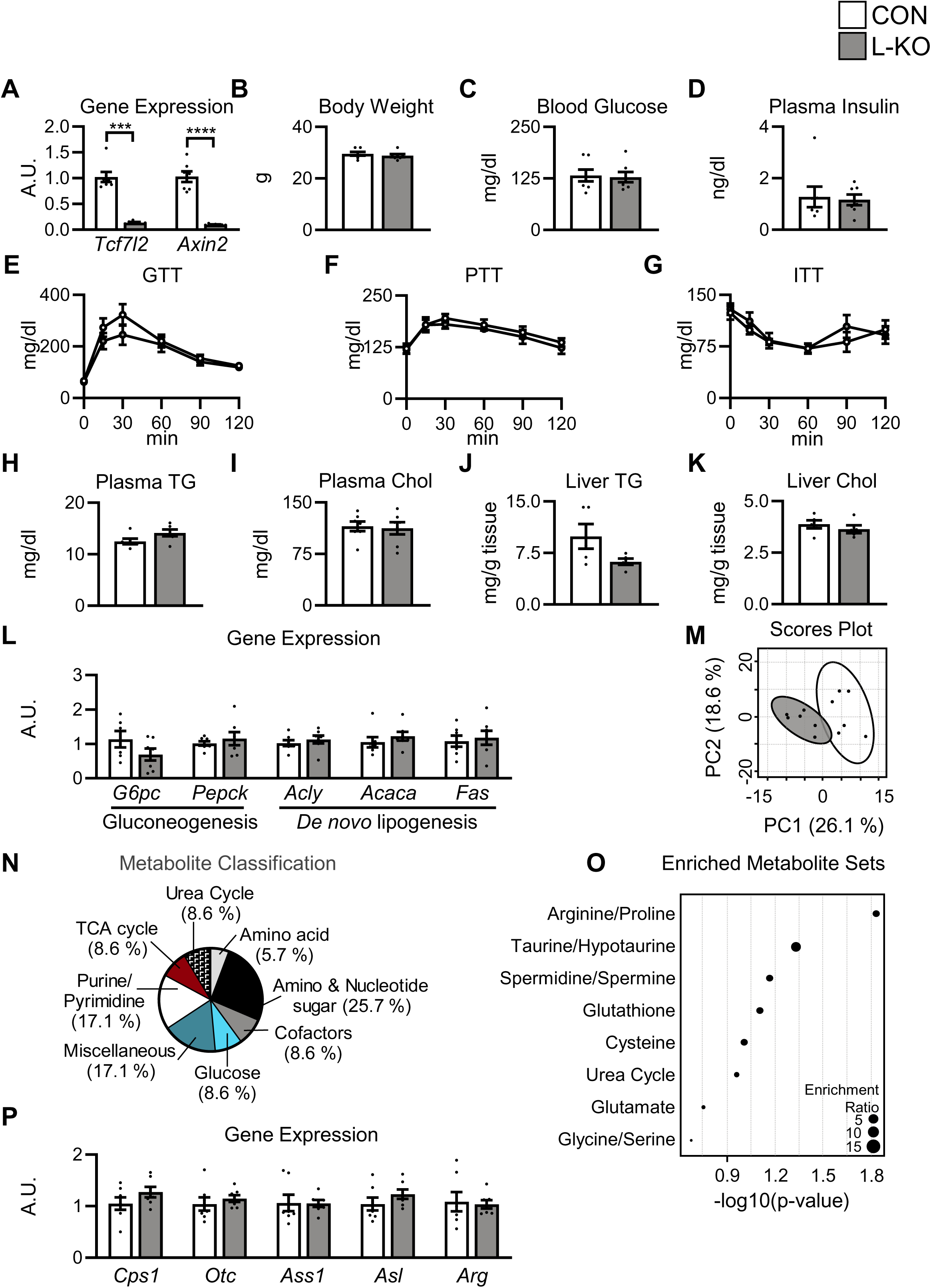
Disruption of hepatic *Tcf7l2* leads to widespread metabolite changes in chow diet-fed male mice. Six to eight week old male *Tcf7l2^Flox/Flox^* mice were injected with adeno-associated virus encoding either GFP (CON) or Cre (L-KO) and placed on a chow diet for twelve weeks. (**A**) Liver QPCR analysis. (**B**) Final body weight. (**C, D**) Four hours fasting (**C**) blood glucose and (**D**) plasma insulin. (**E**) Blood glucose levels during glucose tolerance test (GTT), (**F**) pyruvate tolerance test (PTT), and (**G**) insulin tolerance test (ITT). Four hours fasting (**H**) plasma triglycerides (TG) and (**I**) cholesterol (Chol). (**J**) Liver triglycerides (TG) and (**K**) cholesterol (Chol). (**L**) Liver QPCR analysis. (**M-O**) Liver metabolite profiling. (**M**) Principal Component Analysis (PCA). (**N**) Significantly altered liver metabolites (-log10(pval)>1.3). (**O**) Metabolite Set Enrichment Analysis (MSEA). (**P**) Liver QPCR analysis. Data are presented as the mean ± SEM; n=5-7/group. A.U., arbitrary units.

**Suppl. Figure 2:**
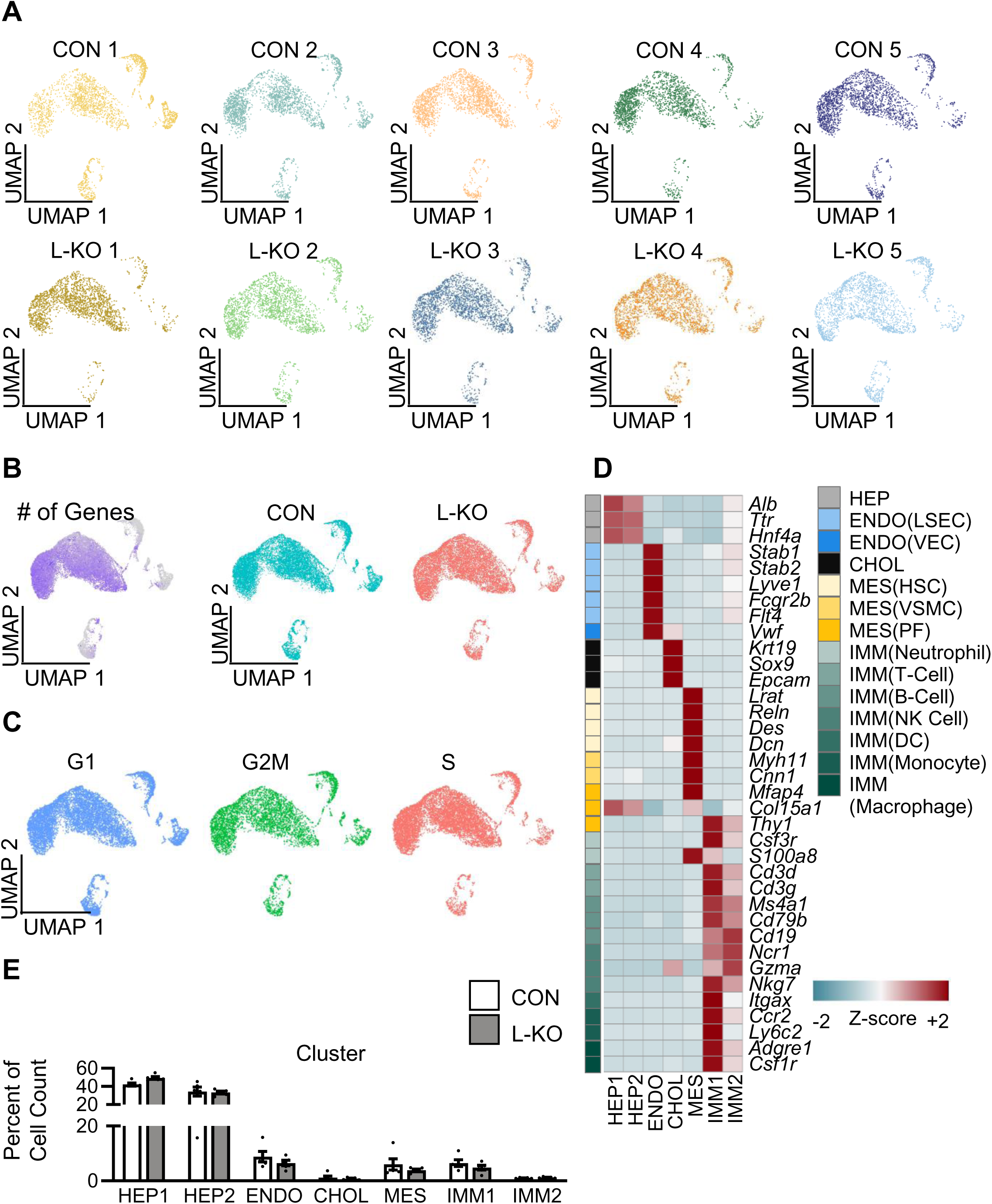
snRNA sequencing quality control. Five to six week old male *Tcf7l2^Flox/Flox^*mice were injected with adeno-associated virus encoding either GFP (CON) or Cre (L-KO) and placed on a chow diet for 12 weeks. UMAP visualization by (**A**) sample; (**B**) number of genes and genotype; and (**C**) cell cycle phase. (**D**) Heatmap of cell type markers. (**E**) Cell count by cluster as a percentage of total by genotype.

**Suppl. Figure 3:**
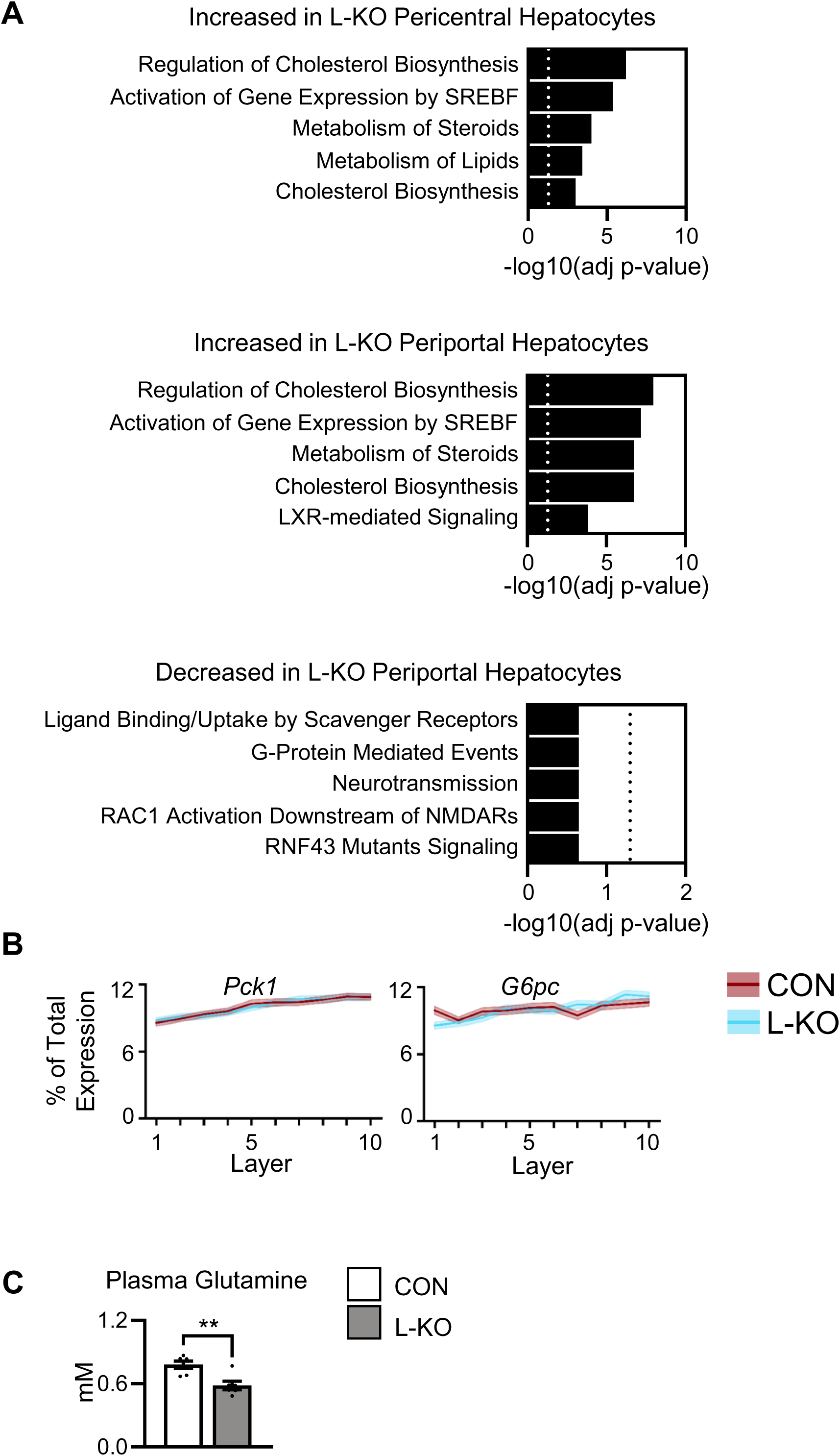
Disruption of hepatic *Tcf7l2* results in changes in pericentral and periportal gene expression. Five to eight week old male *Tcf7l2^Flox/Flox^* mice were injected with adeno-associated virus encoding either GFP (CON) or Cre (L-KO) and placed on a western diet for 12 weeks. (**A**) Genes that were differentially expressed between pericentral (HEP2) or periportal (HEP1) hepatocytes from CON and L-KO livers by snSEQ were subjected to Enrichr analysis as described in methods. Five most significant gene sets are shown. Dotted line marks significance threshold of -log10(adj p-value)>1.3. (**B**) The distribution of gene expression across the lobule using snSEQ analysis. (**C**) Plasma glutamine levels. *P* values were determined by Student’s *t*-test;\*\**P* < 0.01.

**Suppl. Figure 4:**
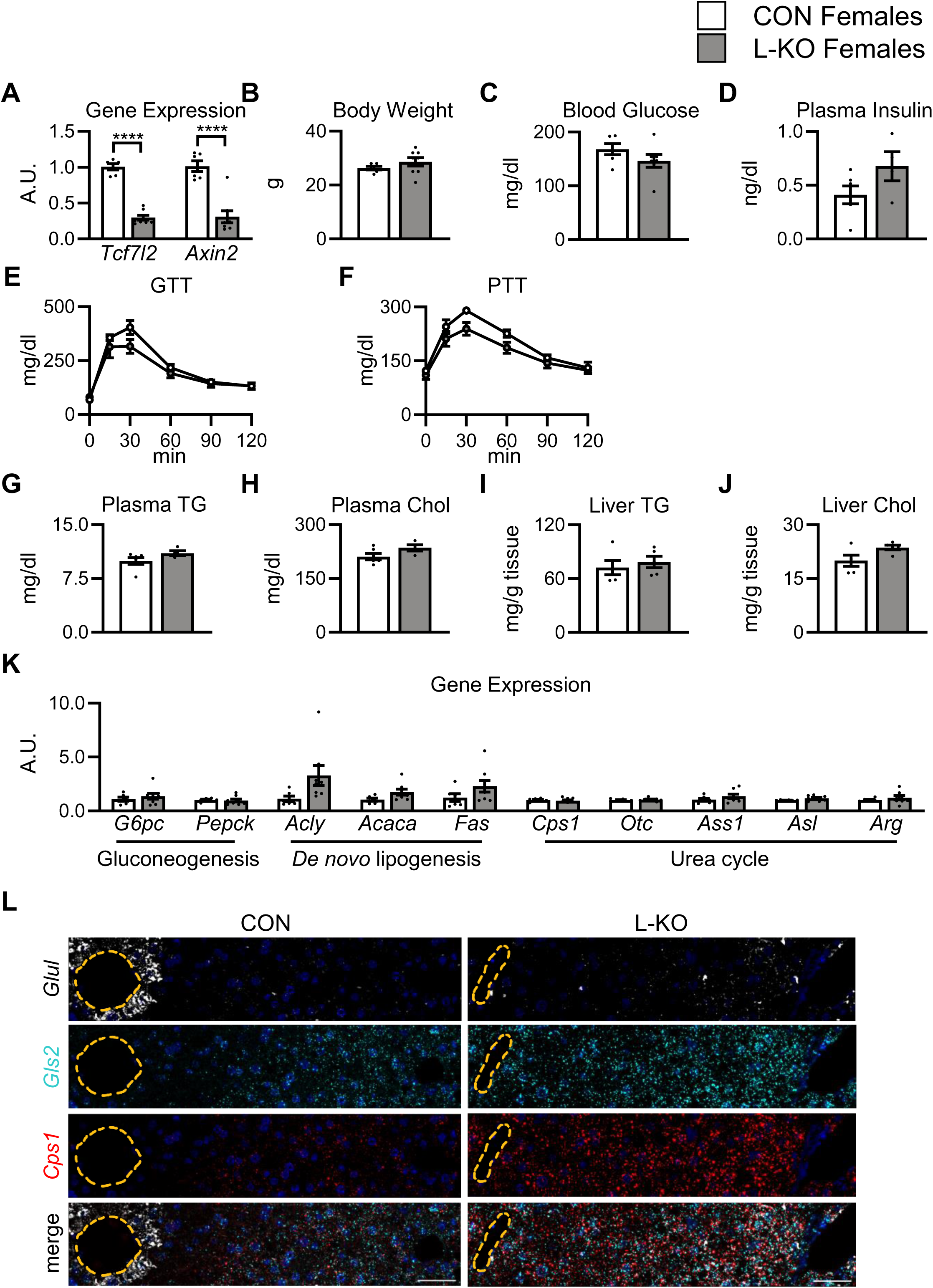
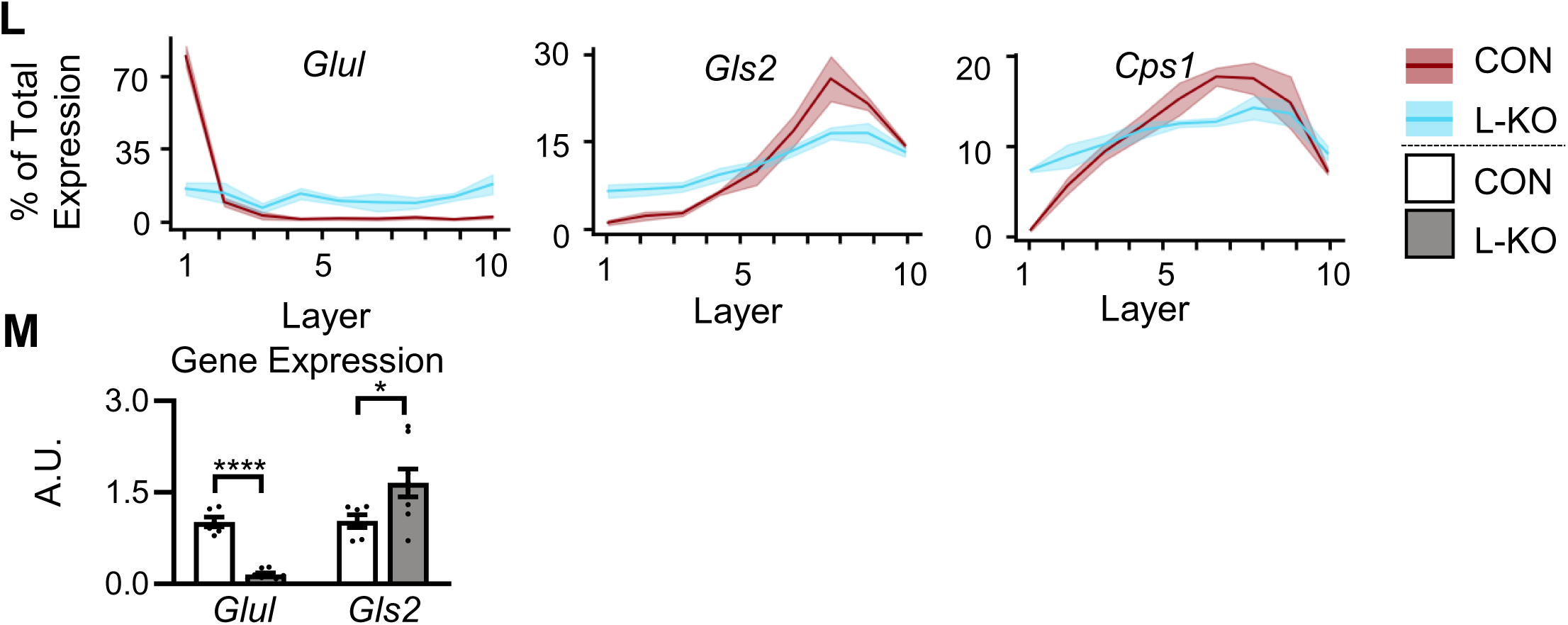
Disruption of hepatic *Tcf7l2* leads to metabolic changes in western diet-fed female mice. Six to eight week old female *Tcf7l2^Flox/Flox^* mice were injected with adeno-associated virus encoding either GFP (CON) or Cre (L-KO) and placed on a western diet for twelve weeks. (**A**) Liver QPCR analysis. (**B**) Final body weight. (**C, D**) Four hours fasting (**C**) blood glucose and (**D**) plasma insulin. (**E**) Blood glucose levels during glucose tolerance test (GTT) and (**F**) pyruvate tolerance test (PTT). Four hours fasting plasma (**G**) triglycerides (TG) and (**H**) cholesterol (Chol). (**I**) Liver triglycerides (TG) and (**J**) cholesterol (Chol). (**K**) Liver QPCR analysis. (**L**) Representative images of smFISH and quantification. Note: RBC were masked out in *Glul* (FITC) channel as they had high fluorescence. (**M**) Liver QPCR analysis. Data are presented as the mean ± SEM; n=4-8/group. *P* values were determined by Student’s *t*-test; \**P* < 0.05, \*\*\*\**P* < 0.0001. Central vein highlighted by dashed orange line; scale bar = 40 µm. A.U., arbitrary units.

